# Inference of Chromosome-length Haplotypes using Genomic Data of Three to Five Single Gametes

**DOI:** 10.1101/361873

**Authors:** Ruidong Li, Han Qu, Jinfeng Chen, Shibo Wang, John M. Chater, Le Zhang, Julong Wei, Yuan-Ming Zhang, Chenwu Xu, Wei-De Zhong, Jianguo Zhu, Jianming Lu, Renyuan Ma, Sergio Pietro Ferrante, Mikeal L. Roose, Zhenyu Jia

## Abstract

Knowledge of chromosome-length haplotypes will not only advance our understanding of the relationship between DNA and phenotypes, but also promote a variety of genetic applications. Here we present *Hapi*, an innovative method for chromosomal haplotype inference using only 3 to 5 gametes. *Hapi* outperformed all existing haploid-based phasing methods in terms of accuracy, reliability, and cost efficiency in both simulated and real gamete datasets. This highly cost-effective phasing method will make large-scale haplotype studies feasible to facilitate human disease studies and plant/animal breeding. In addition, *Hapi* can detect meiotic crossovers in gametes, which has promise in the diagnosis of abnormal recombination activity in human reproductive cells.

## Introduction

A haplotype in a diploid individual is a set of DNA variants on a chromosome that are co-inherited from a parent. Knowledge of haplotypes has an essential role in interpreting personal genomes and guiding individualized treatment plans in precision medicine [1, 2]. Haplotype data have also been utilized in many areas of genetic studies, including imputation of low-frequent variants [3, 4] and characterization of DNA-phenotype associations [5, 6]. Numerous GWAS studies have indicated that while single-SNP analysis is not optimal, joint analysis of multiple SNPs along chromosomes, *i.e.*, haplotypes, showed significantly increased power for detection of genetic determinants for complex traits [7, 8].

Determination of haplotypes, termed phasing or haplotyping, is the process of inferring haplotype architecture based on genotypic data using statistical or bioinformatic approaches. The most widely used haplotyping strategy is to phase common genetic variants using population data [9-16]; however, this approach is incapable of phasing *de novo* mutations, rare variants or structural variants, and is limited to infer short-range haplotype fragments, which constrains its use in genetic studies as well as precision medicine [2]. Experimental approaches targeting whole-chromosome phasing involve the physical separation of homologous chromosomes in diploid cells using chromosome microdissection, FACS-mediated chromosome sorting, or microfluidics, followed by single-chromosome sequencing [17-19]. Nevertheless, these approaches usually require specialized equipment which is expensive and are typically time-consuming. Numerous sequencing technologies including fosmid-based dilution pool sequencing, long fragment read (LFR) technology, PacBio single molecule real-time (SMRT) long-read sequencing, 10X Genomics linked-read sequencing, and proximity ligation (Hi-C) sequencing can also be employed to generate long-range haplotype fragments [20-23]. A novel single-cell DNA template strand sequencing (Strand-seq) technique has been invented to sequence either Watson strand or Crick strand of a chromosome in a diploid somatic cell and phase chromosomal haplotypes using pooled Strand-seq libraries [24, 25]. However, the cost associated with these sequencing technologies are still high, making large-scale research infeasible.

Gamete cells such as pollen grains in plants or sperms and eggs in animals are the natural packaging of haploid complements that are formed by meiotic recombination. Using haploid data of single gamete cells may substantially reduce the complexity in inferring the donor’s chromosomal haplotypes, compared to the phasing approaches using diploid data of somatic cells. However, the development of gamete-based phasing methodologies is still in the early stage, requiring either a large number of gametes or manual inspection for assembly to ensure phasing accuracy [26-28]. No cost-efficient and user-friendly software has been made available for phasing chromosome-length haplotypes with gamete data. To fill this void, we developed an innovative method, named *Hapi* (<u>Hap</u>lotyping with <u>i</u>mperfect genotype data), for a fully-automatic inference of an individual’s chromosomal haplotypes using 3 to 5 gametes, given the heterozygous loci are already known for the genome of this individual. Comprehensive comparisons, involving the use of a simulated dataset, a maize microspore dataset, and a human sperm sequencing dataset, demonstrated that the new *Hapi* method outperformed two existing approaches in terms of phasing accuracy and cost efficiency. The results also suggested that chromosomal haplotypes may be inferred by using only 3 gamete cells if the genotype data are of high quality. Simple, inexpensive and reliable techniques for isolation, lysis, and whole-genome amplification (WGA) of single gamete cells allied with the new *Hapi* method will make the genome-wide haplotype association study (GWHAS) affordable and feasible. In addition, the crossover analysis module in the *Hapi* R package can be used for analysis of crossovers on gamete chromosomes, which will facilitate research on meiotic recombination and also potentially lead to adoption by the public health sector, such as diagnosis of abnormal recombination activity in human sperms and eggs to aid in reducing infant mortality, birth defects, and miscarriages.

## Results

### Implementation of *Hapi*

Implementing the *Hapi* algorithm to phase an entire chromosome consists of three steps: (1) data preprocessing, (2) inference of draft haplotypes, and (3) assembly of high-resolution chromosomal haplotypes (Fig. 1). In step (1), markers with potential genotyping errors in any gamete cells are filtered out *via* an iterative Hidden Markov Model (HMM) analysis of gamete pairs (Supplementary Fig. 1; see **Methods**). A subset of markers, which have been successfully genotyped in at least 3 gametes, are selected to form a “precursor” framework. In the framework, missing data in each gamete are iteratively imputed using supporting data in other gametes (Supplementary Fig. 2; see **Methods**). The markers, usually of a small number, with missing data that cannot be fully resolved by imputation are eliminated, resulting in the final framework for building draft haplotypes. In step (2), the draft haplotypes are derived by sequentially analyzing pairs of adjacent framework markers using the majority voting method, through which the haplotypes for each marker pair are determined by the link type represented in the majority of the gametes (Supplementary Fig. 3; see **Methods**). The maximum parsimony of recombination (MPR) principle is then adopted to proofread disputable positions of the draft haplotypes (Supplementary Fig. 4; see **Methods**). In step (3), each gamete chromosome is compared to the draft haplotypes to identify haplotype-converting points (HCPs) to deduce gamete-specific haplotypes, with the non-framework markers being phased. Consensus high-resolution haplotypes are eventually determined from these gamete-specific haplotypes by voting for the major allele at each locus (Supplementary Fig. 5; see **Methods**).

**Fig. 1:**
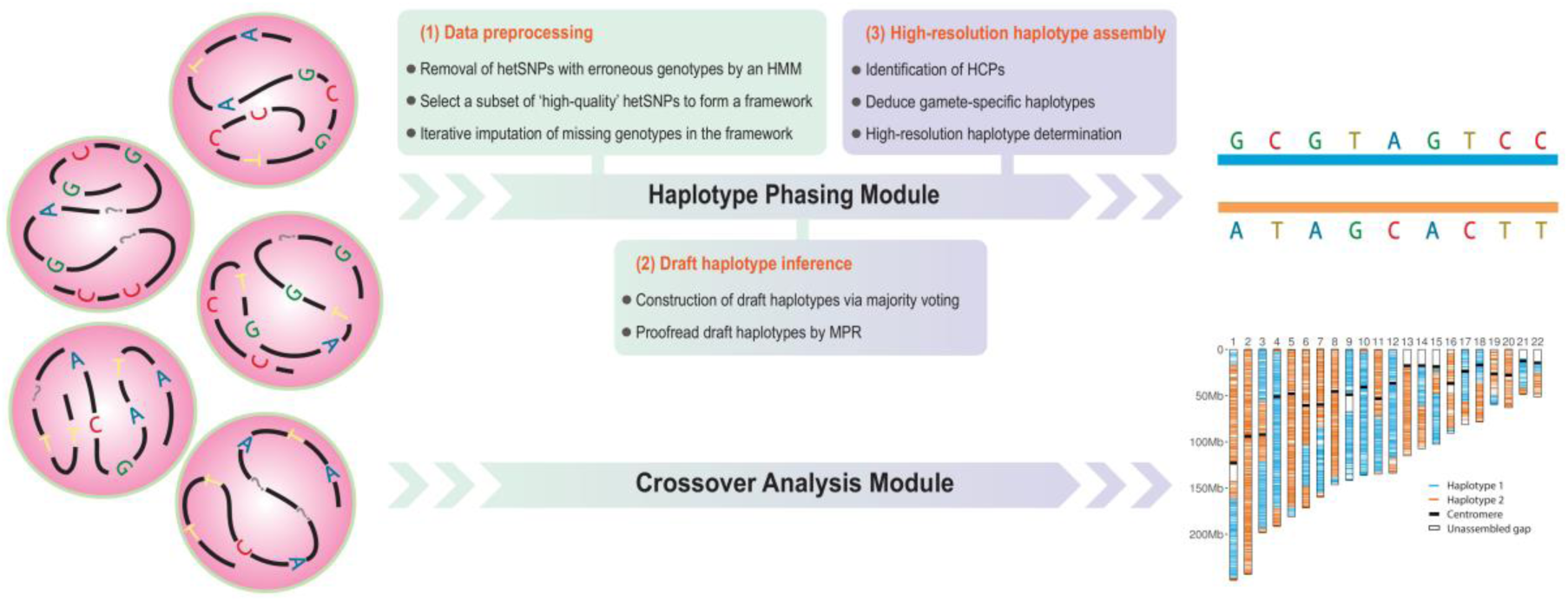
Flowchart of the *Hapi* phasing pipeline.

A user-friendly R package has been developed for implementing the *Hapi* algorithm to infer chromosome-length haplotypes using single gamete cells. *Hapi* uses genotype data of hetSNPs in individual gametes and outputs the high-resolution chromosomal haplotypes as well as confidence level of each phased hetSNP. The package also includes a crossover analysis module allowing downstream analyses and visualization of crossover positions identified in the observed gametes (Fig. 1). The *Hapi* package is publicly available at https://github.com/Jialab-UCR/Hapi.

### Comparison of phasing methods in a simulated dataset

We carried out a comprehensive simulation study to compare the performance of *Hapi* with the other two competitive methods, *One*-*Versus*-*All* (*OVA)* [28] and *Pairwise HMM (PHMM)* [26]. Three factors that may affect phasing accuracy and cost efficiency were considered in each scenario, *i.e.*, (1) the number of hetSNP markers on the chromosome, (2) the number of gametes, and (3) the frequency of missing genotype data. In the simulated dataset, a pool of 100 haploid gametes was generated from a single diploid donor. The number of hetSNPs on the chromosome ranged from 5,000 to 1,000,000. Three to fifteen gametes, each with 0 to 3 crossovers on the chromosome, were arbitrarily selected from the 100 haploid gametes without replacement. 10% to 70% of missing genotype data were randomly introduced to each simulated gamete chromosome. Moreover, 1% genotyping errors were randomly placed on the simulated gamete chromosomes. We compared the three methods under different scenarios with predetermined number of gametes, number of hetSNPs, and missing genotype rate. 100 replicates were performed for each scenario. A successful inference was defined if more than 99% of hetSNPs were correctly phased in each replicate run.

The results indicated that *Hapi* outperformed the other two methods in both phasing accuracy and cost efficiency (Fig. 2). When 5,000 hetSNPs per chromosome were considered, *Hapi* only needed 6 gametes to correctly infer haplotypes even with 60% of missing genotype data. For *OVA*, at the missing rate of 50%, the first 100% correct inference of haplotypes occurred when 7 gametes were used. However, when more gametes were included in the analysis, the performance of *OVA* was not monotonically increased, indicating a lack of reliability and robustness of the method. If 70% of the marker data were missing, *Hapi* was able to reconstruct haplotypes correctly with 11 or more gametes; whereas, *OVA* failed to do so even when all 15 gametes were used. With increased density of hetSNPs, fewer gametes were needed and a higher rate of missing genotypes can be tolerated for both methods to correctly phase the chromosome, however, *Hapi* always outcompeted *OVA* by requiring fewer gametes and allowing more missing data. The results also indicated that only 3 gametes may be enough for successful inference of chromosomal haplotypes when gamete data are of high quality. *PHMM* behaved quite differently from the other two methods. The performance of *PHMM* did not change with the rate of missing data, while the performance was barely improved with the increase in number of hetSNPs. Rather, the phasing accuracy of *PHMM* depended on the number of gametes used in analysis. In general, many more gametes are required for *PHMM* to infer correct haplotypes than the other two methods.

**Fig. 2:**
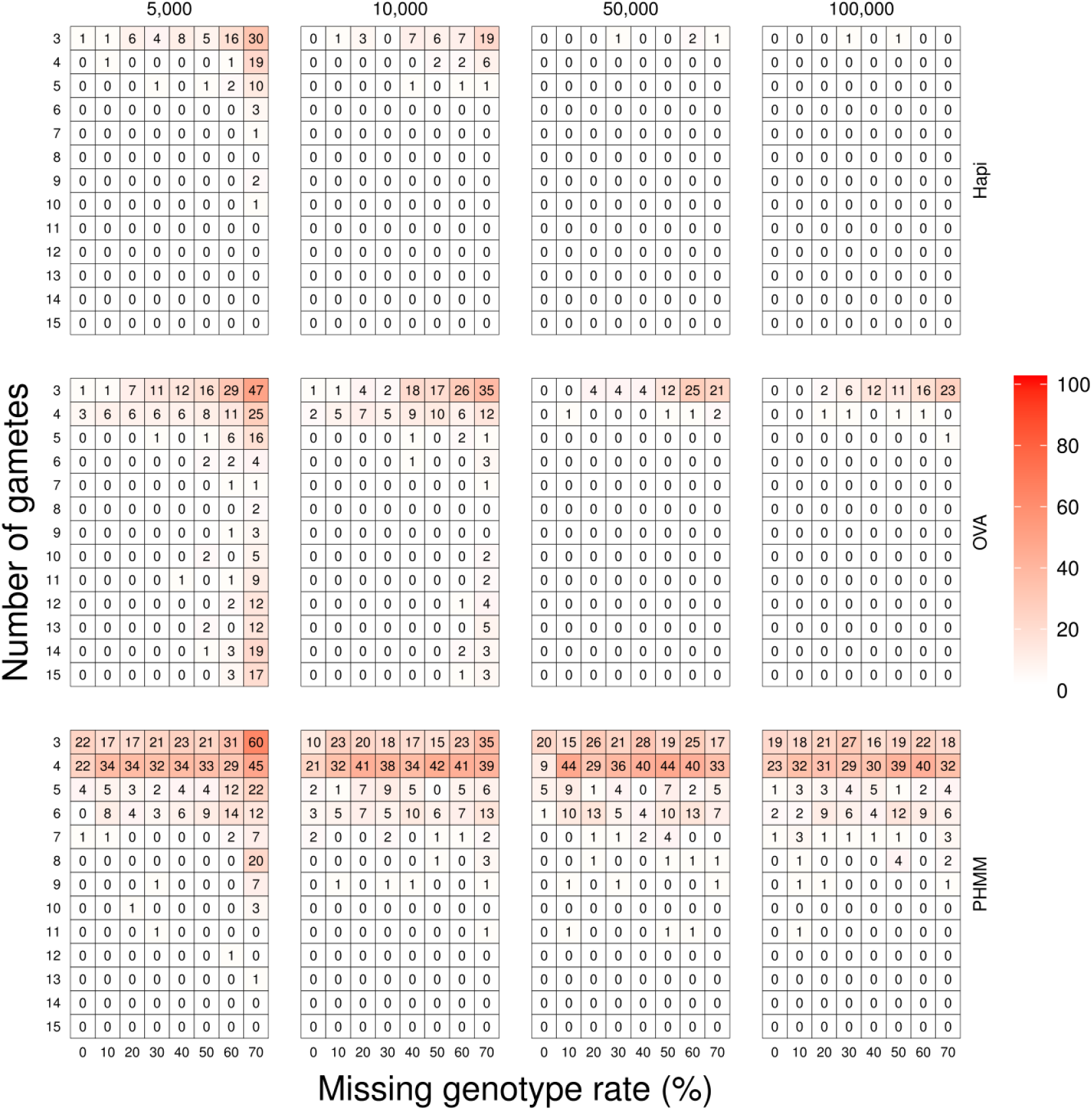
Performances of three methods (*Hapi*, *OVA*, and *PHMM*) in the simulated dataset for 5,000 to 100,000 hetSNPs per chromosome. The number in each heatmap grid denotes for how many times out of the 100 replicates the haplotypes are incorrectly inferred in that scenario.

### Comparison of phasing methods in the maize microspore dataset

A maize microspore sequencing dataset from F1 hybrid individuals of a cross between two inbred lines [29] was used to further evaluate the performance of the three methods. This is an ideal validation dataset since the parental haplotypes are known. To avoid using microspores from the same meiosis event, one microspore from each of the 24 tetrads was randomly selected to form a 24-gamete pool. The number of hetSNPs on the maize chromosomes ranges from 42691 (Chr10) to 82689 (Chr1). The average rate of missing genotype data for 10 chromosomes across the 24 selected gametes is about 50%, with the maximum missing rate equal to 72.46% (Supplementary Table 1). To phase a maize chromosome, the 24 selected gametes were sorted in descending order of missing rates on that chromosome, *i.e.*, the first gamete in the sorted list has the most missing data for the chromosome. 3 to 15 gametes were sequentially selected from the sorted list and analyzed with the three methods, respectively, to infer haplotypes for that chromosome. This process was repeated to phase all 10 chromosomes, yielding a total of 390 scenarios (13 numbers of gametes × 10 chromosomes × 3 methods). In each scenario, the phased chromosome was compared with the known parental haplotypes to calculate phasing accuracy. A successful inference of chromosomal haplotypes is defined if > 99% of the markers were correctly phased.

The results indicated that the *Hapi* method can achieve phasing accuracies of greater than 99.9% in most scenarios, with two exceptions at 98.46% for Chr2, and 99.89% for Chr6, respectively. The accuracy lower than 99% happened when only 3 gametes were analyzed for phasing Chr2 (Fig. 3). A close look at Chr2 of these 3 gametes disclosed two crossovers on two gamete chromosomes in a small region (39 hetSNPs in between) near one end of the chromosome. In order to construct a reliable draft haplotype, *Hapi*, by default, excludes any small block (< 100 hetSNPs) delimited by two close-in crossovers from the draft haplotypes, prior to implementation of MPR; thus, in some cases, the phase of the two merging framework markers may be incorrectly inferred by misinterpreting the link types in between due to the removed two crossovers. On the other hand, at least 6 and 7 gametes are required for *OVA* and *PHMM*, respectively, to achieve a phasing accuracy of > 99% for all the 10 chromosomes.

**Fig. 3:**
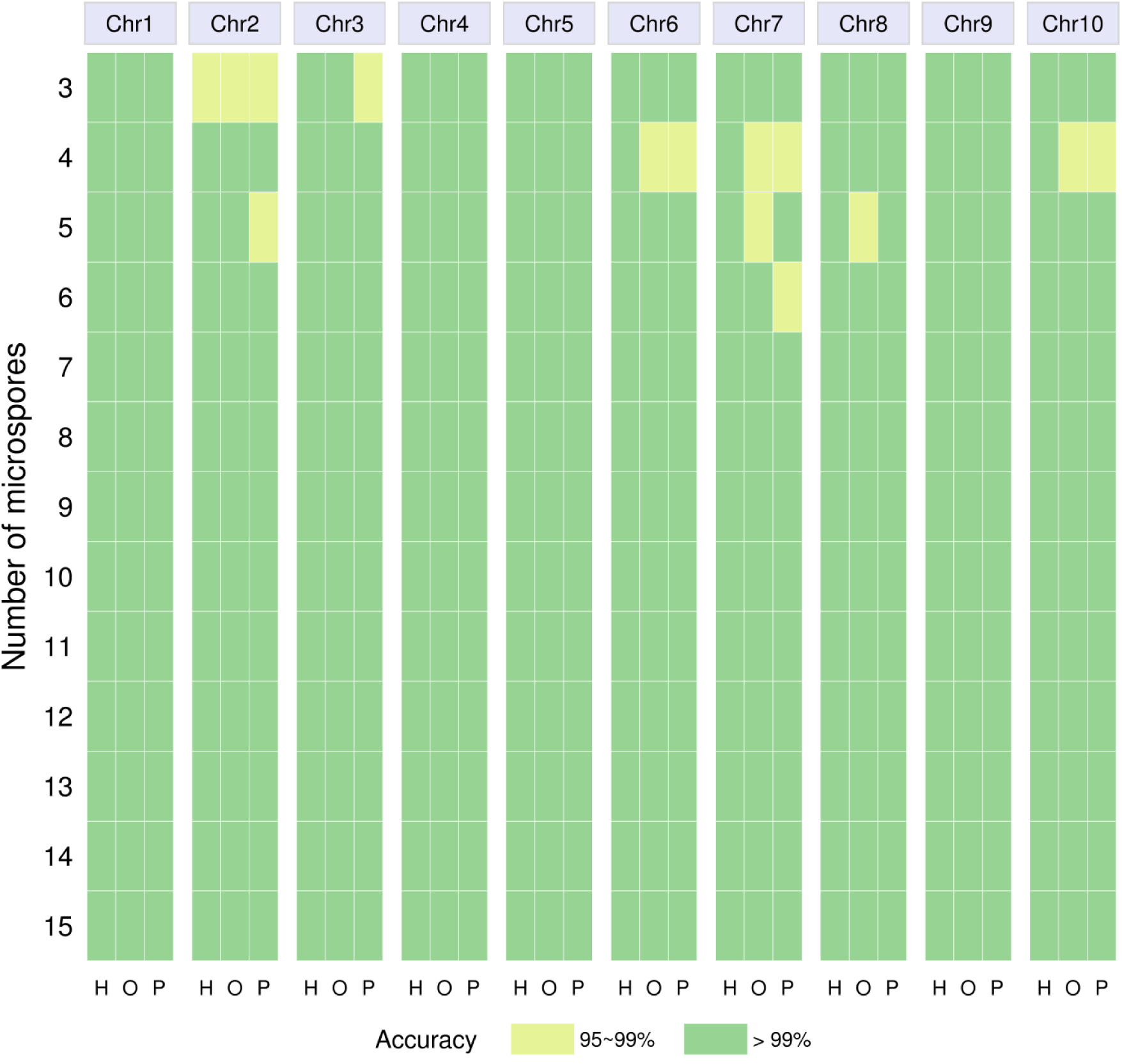
Performances of three methods (H: *Hapi*, O: *OVA*, and P: *PHMM*) on phasing 10 chromosomes of maize in the maize microspore sequencing dataset.

### Comparison of phasing methods in the human sperm dataset

To further benchmark the three phasing methods, a human sperm sequencing dataset consisting of 11 independent sperm cells from the donor of the HuRef diploid genome sequence was used [28]. Although the true haplotypes for this donor are unknown, a ‘phased’ genome consisting of 1.82 million hetSNPs has been suggested based on a joint analysis of these 11 sperms sequenced at 1.5~3.7× coverage and 16 additional sperms genotyped using the Illumina HumanOmni-Quad v1.0 BeadChip (array data not publicly available) [28]. The raw sequencing data of the 11 sperm cells were downloaded and 1.66 million out of the 1.82 million hetSNPs were called in at least one sperm. The number of hetSNPs on 22 autosomes ranges from 15340 (Chr22) to 141669 (Chr2), and the rate of missing genotype data ranges from 70.95% to 86.49% (Supplementary Table 2). When phasing a human chromosome, the 11 sperms were sorted in a similar manner as for the maize data based on the missing genotype rates. 3 to 11 sperms were sequentially selected from the sorted list and analyzed using the three methods described above to infer chromosomal haplotypes which were then compared with the hetSNPs ‘phased’ in the original study[28] to calculate the concordance rate. Since the chromosomal haplotypes suggested in the original study may be subject to errors, we relaxed the criterion in the sperm analysis by defining a successful inference of haplotypes if > 95% of phased markers are in agreement with the haplotypes suggested by Kirkness et al [28].

The results showed that *Hapi* can correctly phase all 22 autosomes with 3 sperms; whereas, *OVA* and *PHMM* required at least 7 and 8 sperms, respectively, to achieve the same level of accuracy (Fig. 4). When 7 or less sperms were used, *Hapi* performed consistently well but the performances of *OVA* and *PHMM* fluctuated wildly, indicating *Hapi* provides more reliable phasing results with small samples. Interestingly, *PHMM* can correctly infer the haplotypes of chromosome 1 with 6 to 10 gametes but failed when all 11 sperms had been used. Although a consistency of 95% was used to determine the success of haplotype inference, *Hapi* achieved > 99% of consistency for 82% of the scenarios (164 out of 198). For *Hapi*, the majority of scenarios with consistencies of 95%~99% were for the analyses of Chr15, Chr16, and Chr21, which also appeared to be challenging to the other two approaches, suggesting a complication in the genotype data for these chromosomes. Overall, among the 1.66 million hetSNPs phased by *Hapi* using all the 11 sperms, 99.73% (1,658,197/1,662,611) of them are concordant with the haplotypes suggested by Kirkness et al [28]. An inspection of the non-concordant hetSNPs showed that 49.1% of them are only supported by 1 sperm and 33.4% of them have discordancy among 2 or more supporting sperms. The disputably phased hetSNPs tend to cluster around the centromere or at either end of the chromosomes (Supplementary Fig. 6). The hetSNPs that are not in agreement between *Hapi* and the suggested haplotypes on Chr15 are evenly distributed along the chromosome, which might be ascribed to the complication in data of sperm Y47 being contaminated by DNA from other lysed cells as mentioned in the original paper [28]. The results showed that phasing Chr15 is equally challenging for *OVA* and *PHMM*. Compared with *Hapi*, the major deficiency in haplotype phasing with *OVA* and *PHMM* is due to their core strategy of a direct inference of crossover positions, which is sensitive to the regions with ambiguous genotypes or complications caused by multiple crossovers in more than one gamete. For example, as shown in the Supplementary Fig. 7, if 10 sperms were analyzed, a crossover on Chr1 in the sperm X69 (reference chromosome) was not claimed because it was only supported by 5 out of 9 other sperms and missed the cutoff of ≥ 0.6 for determining a crossover. However, when including the 11th sperm, the crossover became supported by 6 out of 10 sperms, which claimed a false crossover and yielded an incorrect gamete-specific haplotype. In *Hapi*, such genomic regions harboring complicated multiple cv-links will be excluded from the draft haplotypes to reduce the chance of phasing errors. In addition, a special capping function has been designed in *Hapi* to phase either end of the chromosome, which are usually excluded from the framework but may also involve recombination. The *OVA* method, which also leverages draft haplotypes for phasing a chromosome, cannot handle recombination beyond the framework.

**Fig. 4:**
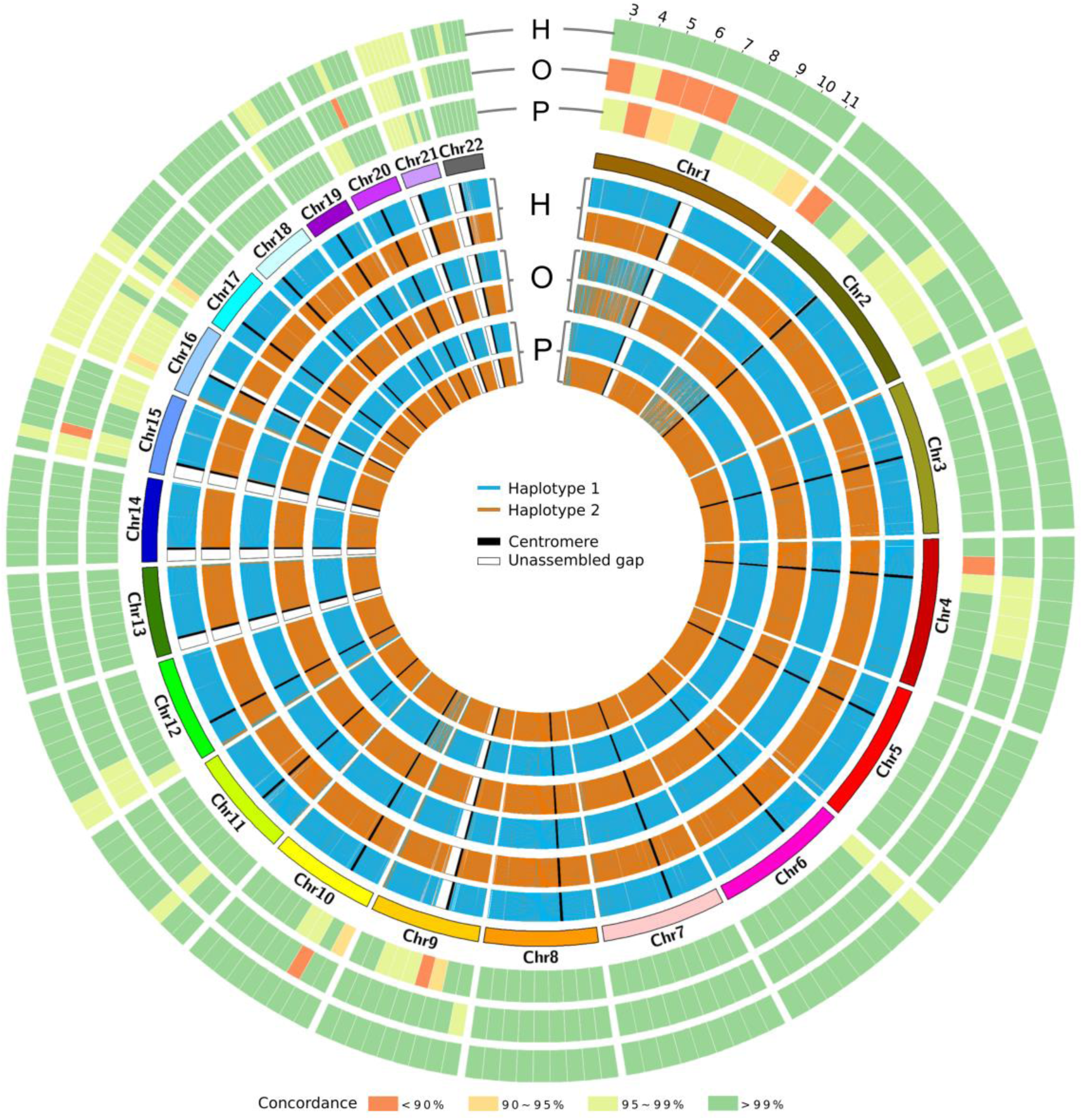
Performances of three methods (H: *Hapi*, O: *OVA* and P: *PHMM*) on phasing 22 autosomes in the human sperm dataset. The 3 outer circles show the phasing concordance with the suggested haplotypes [28] for each method using 3 to 11 sperms. The 6 inner circles are the haplotypes inferred by the 3 methods using 3 sperms with the most missing genotypes.

### Recombination analysis in the human sperm dataset

With the phased chromosome-length haplotypes, an HMM was used to infer crossover positions in the sperm genomes by successively contrasting hetSNPs in each sperm with the inferred chromosomal haplotypes (Supplementary Fig. 8).

A total of 254 crossovers along the 22 autosomes were identified in the 11 sperms with an average of 1.05 per chromosome. Compared with the 260 crossovers identified in the original paper [28], 251 were also identified by the *Hapi* method (Supplementary Table 3). The 12 inconsistent crossovers are all located at the ends of chromosomes, and such inconsistency may be ascribed to either of two following reasons. (1) In general, the *OVA* method in the original paper cannot accurately infer haplotypes at the chromosome ends, yielding incorrect crossovers in those regions. (2) The observed double crossovers in a very small region are considered to be either caused by a gene conversion event or consecutive genotyping errors and thus are filtered out by *Hapi.* The number of crossovers was counted in each bin (5Mb in length) along 22 autosomes and distributions of the 254 crossovers are depicted in Fig. 5A. The resolution of crossover locations ranges from 79bp~788kb with a median of 89.3kb, which is roughly the same as the 82.5kb resolution reported in the original paper [28]. Over 75% of the 254 crossovers were located within an interval of < 200kb (Fig. 5B). Distribution of distances between any two chromosomally adjacent crossovers was provided (Fig. 5C), which can be used for recombination-relevant research including location of hot spots or interference in the formation of chromosomal crossovers during meiosis. Functions for downstream analysis and visualization are included in the ‘crossover analysis’ module of the *Hapi* package.

**Fig. 5:**
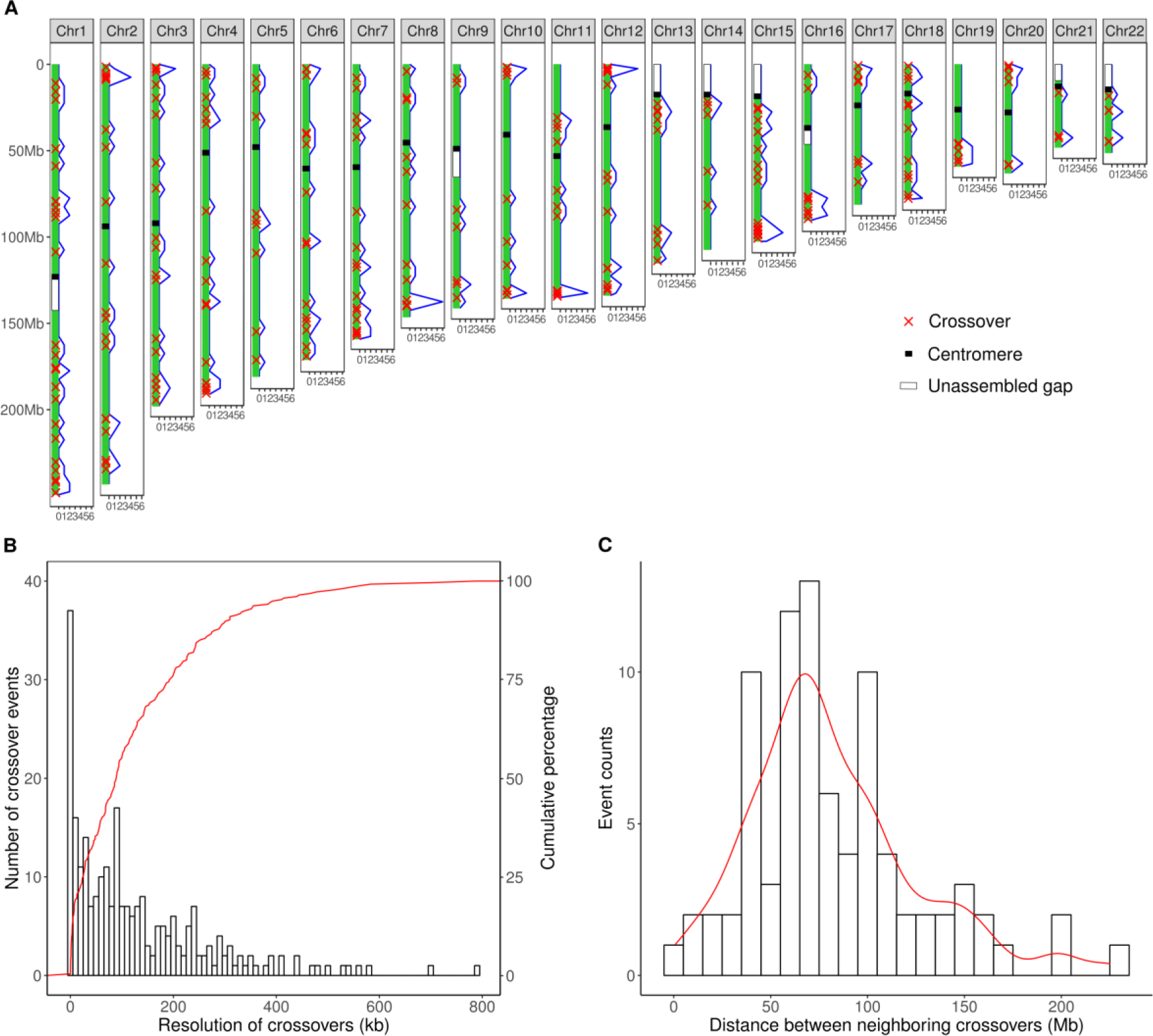
Crossover analysis in the human sperm sequencing dataset. (**A**) The distribution of 254 identified crossovers on the 22 autosomes. (**B**) The distribution of the crossover resolutions (distance between two adjacent markers that involve a crossover). (**C**) The distribution of distances between two neighboring crossovers.

## Discussion

Current diploid-based haplotyping methods are costly or only produce haplotype fragments, whereas, haploid-based alternatives using gamete data may break through such boundary to infer chromosome-scale haplotypes for individual genomes. Two haploid-based phasing methods both rely on the accurate detection of crossover positions on gamete chromosomes, which may be challenging for complex chromosomal regions with many repetitive DNA elements, such as large segmental duplications. The existence of missing and ambiguous genotype calls makes the task even harder, leading to inaccurately phased haplotypes. In this study, we developed a highly efficient algorithm that only requires 3 to 5 gametes to correctly reconstruct high-resolution chromosomal haplotypes.

Using simulated and real sequencing datasets, we demonstrated that *Hapi* outperforms the other two methods in phasing accuracy, reliability, and cost efficiency. To achieve the same level of phasing accuracy, *Hapi* required fewer gametes and can tolerate more missing genotypes than the other two methods. This is not only because of the sophisticated and improved phasing strategy, but also due to the novel algorithms for handling imperfect data (missing and erroneous genotypes). When different numbers of gametes were used for phasing, *Hapi* performed consistently well but the performances of *OVA* and *PHMM* fluctuated wildly, indicating the new *Hapi* method handles ambiguous data from a small number of gametes very well and produces reliable phasing results. Supplementary Fig. 9 provides an example of showing deficiencies for *OVA* and *PHMM* when 4 gametes are analyzed. If two crossovers, one in a gamete and the other one in another gamete, are located in a 1Mb region, *OVA* cannot detect any crossover while *PHMM* detects 4 crossovers, suggesting that neither method is capable of phasing chromosomes when multiple crossovers occur in multiple gametes within a very small chromosomal region (such as a recombinational “hotspot”). In contrast, this challenge can be well resolved by the majority voting and MPR strategies implemented in the *Hapi* method. Since ambiguous genotypic data sometimes occurs at the chromosome ends, a special capping algorithm has been designed in the *Hapi* method to polish the haplotypes in those regions.

Our study also indicated that 3 gametes may be enough to reconstruct chromosome-length haplotypes by *Hapi* if the genotype data are of high quality, *i.e.*, with few missing or erroneous data. It should be noted that using 3 gametes may fail in a special scenario when two sampled gametes each have a crossover within a very small region. This is because, in the step of proofreading draft haplotypes, small blocks with little genotype information are excluded from the draft haplotypes by default, assuming the probability of having multiple crossovers within these blocks in more than one gamete is low. In this specific but rare scenario, removal of such blocks may lead to the wrong determination of the major link type and thereafter the haplotypes. If only 3 gametes are available, it is recommended to implement the *Hapi* method with and without removing blocks in constructing draft haplotypes and check the consistency in results from two different settings.

Unlike existing phasing algorithms that demand sequencing long-reads or linked-reads in diploid cells, the *Hapi* method can analyze hetSNPs data of single gamete cells generated using any genotyping platform. Either nucleobases (A/T/C/G) or binary code (0/1) can be used as the input genotypic data for hetSNPs in gamete genomes. Advanced technologies, such as 10X Genomics linked-read sequencing, are not necessary for the *Hapi* method, but may be used as ancillary approaches to generate designated long-range haplotype fragments for complex and challenging genomic regions, further perfecting the chromosomal haplotypes inferred by the *Hapi* method.

As a cost-effective method for inferring chromosome-length haplotypes of individual genomes, the new *Hapi* method has made GWHAS feasible and affordable, which will inspire innovative ideas and advance our understanding of the relationship between DNA and phenotypes. Another important application of the *Hapi* package is to implement the crossover analysis module to derive maps of recombination in gametes based on the inferred chromosome-length haplotypes, which will facilitate recombination-relevant research in humans and may be translationally applied in clinical labs to manage human diseases that are associated with abnormal recombination. This unique function can also be used to monitor the crossovers on plant genomes to facilitate more rapid introgression of target genes or to break up undesirable linkages for crop improvement.

## Materials and Methods

### Key Component Algorithms Employed in *Hapi*

#### 1. HMM for detection of genotyping errors

Enlightened by a previous study [26], a HMM is adopted to linearly scrutinize hetSNP markers along the chromosome in two gametes to identify markers bearing genotyping errors (Supplementary Fig. 1). In the HMM, there are two observations ‘s’ and ‘d’ indicating the two possible outcomes, either same or different, in terms of the relationship of observed genotype calls at a hetSNP locus between two gametes. Two hidden states, ‘S’ and ‘D’, represent the invisible relationship between the true genotypes of this marker in these two gametes, with ‘S’ and ‘D’ denoting the same and different genotypes, respectively. The initial probabilities of the two states are 0.5. Because the observed genotype outcomes may be different from the hidden states due to the genotyping errors at rate *E*, the emission probabilities to observe the same genotype calls, *i.e.*, s, given the S hidden state is 1-2*E*(1-*E*) and to observe the different genotype calls, *i.e.*, d, is 2*E*(1*E*). The emission probabilities given the D state are defined in the same way. A transition is defined as a change in state when scanning two adjacent markers, indicating that a meiotic recombination likely occurs between these two markers on either gamete chromosome. Suppose the recombination frequency is *R*, the transition probabilities from one state to itself is 1 - 2*R* × (1 - *R*), and to the other state is 2*R* × (1-*R*). After defining the HMM, Viterbi’s algorithm [30] can be used to determine the most likely hidden state for each marker. Markers with genotyping errors are determined where there are conflicts between the observed outcomes and the inferred states. The HMM is iteratively applied to all gamete pairs for the detection of disputable SNP loci with potential genotyping errors.

#### 2. Imputation of missing genotypes

We define a *framework* as a set of selected hetSNPs for constructing draft haplotypes for each chromosome. Missing data for the framework markers in the gametes are imputed in an iterative manner (Supplementary Fig. 2). When a missing region (either a single marker or consecutive markers) of a ‘target’ gamete is to be imputed, the two markers immediately around this region, called comparator markers, are first compared with those in other ‘support’ gametes. The missing region can be imputed with the information from a support gamete cell only if the genotype calls for these two comparator markers in the target gamete are either both identical or both complementary to those in the support gamete. For example, if genotype calls of the two comparator markers in the target gamete are both identical to those in the support gamete, the missing region on the target gamete is simply imputed with genotype calls of markers in the same region in the support gamete. Otherwise, the missing region in the target gamete is imputed with the reciprocal genotypes in the support gamete. Missing genotypes in one gamete can be eventually resolved only if the imputations are supported by more than 2 support gametes and no imputation conflict is incurred. Once all the gametes are imputed in one iteration, genotypes in the missing regions are updated and the entire process described above will be repeated until no more missing data can be further imputed.

#### 3. Majority voting

With the assumption that recombination is generally rare on the chromosome and even rarer between two neighboring framework markers (a small region) in multiple gametes, the haplotypes of these two adjacent framework markers are deduced by analyzing genotype links (genotype patterns for these two markers) across all gametes based on the majority voting principle. There are two types of links between these two neighboring framework markers, *i.e.*, type I links include genotype patterns 0-0 and 1-1 and type II links include genotype patterns 0-1 and 1-0, where 1 and 0 represent two complementary genotype calls that are arbitrarily and independently assigned at either locus (Supplementary Fig. 3). The most frequent link type is determined as hap-link which represents the likely haplotypes for the two framework markers, whereas the minority link type is considered as cv-link arising from a crossover. The final draft haplotypes can be deduced through walking and voting along the framework of the chromosome.

#### 4. Maximum parsimony of recombination

Maximum parsimony of recombination (MPR) [31], an optimality criterion to search for the haplotype arrangement with minimum number of crossovers in a chromosomal region across all gametes, is adopted by *Hapi* to proofread the equivocal regions (two adjacent framework markers) of draft haplotypes where disputable cv-links have been observed. When five or more gametes are analyzed, we treat any two adjacent markers with 2 or more cv-links as candidate regions for proofreading (Supplementary Fig. 4). If very few (*e.g.*, 3 or 4) gametes are in use, every two adjacent markers with any cv-link are subject to proofreading. The draft haplotypes are first segmented into blocks by the equivocal regions. Small blocks (< 100 hetSNPs) with little genotypic data are excluded from the construction of the draft haplotypes. To phase two neighboring blocks, raw genotype calls (with possible missing data) of the joining hetSNPs markers, *i.e.*, the last 100 consecutive hetSNPs in the first block and the first 100 consecutive hetSNPs in the second block, are retrieved. Since haplotypes within each block are unambiguous, there are only two possible combining haplotypes for these two blocks. The total number of crossovers in all gametes are counted given the two combining haplotypes, and the one generating less crossovers is preferred by the MPR algorithm.

#### 5. Assembly of consensus chromosome-length haplotypes

We arbitrarily select one of the inferred draft haplotypes and use it as a blueprint to deduce gamete-specific haplotypes and eventually assemble the chromosome-length consensus haplotypes through three steps (Supplementary Fig. 5). In step 1, genotype calls of framework markers in each gamete chromosome are compared to the blueprint to identify haplotype-converting points (HCPs) which are caused by potential recombination. These HCPs partition each gamete chromosome into *k* haplotype segments, where *k*-1 is the number of HCPs identified for this gamete chromosome. For the segments 1 through *k*, genotype calls of hetSNPs in every second segment are flipped to form a gamete-specific haplotype, where ‘flip’ refers to switching the current genotype call to its reciprocal genotype. In step 2, each gamete-specific haplotype is synchronized with the blueprint by either remaining the same or flipping over the genotypes of entire chromosomal hetSNPs. In step 3, the first consensus chromosome-length haplotype is reconstructed *via* voting for the most frequent allele at each hetSNP locus across all the gamete-specific haplotypes. The second consensus haplotype is obtained by simply flipping genotypes of hetSNPs on the first chromosome-length haplotype.

If a crossover occurs at the end of a gamete chromosome where hetSNPs are not enclosed in the framework, it becomes challenging to correctly infer the haplotypes for this chromosome-tip region. *Hapi* employs an additional capping strategy to polish two ends of chromosomal haplotypes. First, hetSNPs in such a region are combined with the immediately adjacent 200 consecutive hetSNPs at the joining end of the framework to form a capping block, of which the haplotypes can be inferred by treating them as a small chromosome. Then, small-scale draft haplotypes are constructed for the selected framework markers of this capping block by using the most frequently represented genotype calls across the gametes. The same strategy is adopted to generate gamete-specific haplotypes to deduce consensus haplotypes for this small chromosome-tip region. Lastly, the inferred haplotypes for the capping block are integrated into the chromosome-length haplotypes to accomplish the assembly.

### Rival Phasing Methods

#### 1. *One*-*versus*-*All* (*OVA*) pipeline

Kirkness et al. [28] proposed a two-stage strategy to infer chromosome-scale haplotypes by combining the use of genotyping array data and next-generation sequencing data of sperm cells. In the first stage, array data with relatively high call rate (50.9% on average in their study) were analyzed in a one-versus-all fashion to identify crossovers in the gametes, which were used to construct the draft haplotypes. When phasing a chromosome, a gamete is first set as a reference, and the other gametes are considered as offspring. HCPs are identified for all reference-offspring pairs, where an HCP indicates the position with a potential crossover either on the reference or on offspring chromosome. A crossover is assigned to the reference chromosome if the HCP is identified in the majority of reference-offspring pairs, for example, 13 out of 15 pairs as indicated in the original paper [28]. Otherwise, multiple crossovers must have taken place on the offspring chromosomes. A manual inspection step is required to confirm the crossover locations on each reference chromosome. As a result, a gamete-specific chromosome-scale haplotype can be inferred by the crossovers assigned to the reference chromosome. The entire process described above is repeated until each gamete has been set as a reference for one time. Draft haplotypes can be constructed using these gamete-specific haplotypes by voting for the major allele at each locus. In the second stage, the inferred crossover positions are employed again to assist the analysis of the additional sequencing data of sperm cells to infer the high-resolution consensus chromosome-scale haplotypes.

To perform the comparison analysis, this algorithm was written in R language by us with a few optimizations. (1) Rather than using two sets of gamete genotype data, *i.e.*, SNP array data and sequencing data as in the original study [28], only one dataset is used for the modified two-stage *OVA* pipeline. (2) The gamete genotype data are preprocessed to remove markers with potential genotyping errors and a subset of high-quality hetSNPs is selected to infer crossovers in the gametes. (3) A HMM is used to detect HCPs with higher level of accuracy. (4) A well-written R function is developed to automatically determine crossovers, which gives the exact results of locating recombination, to replace the manual inspection required in the original pipeline.

#### 2. Pairwise HMM (*PHMM*)

The *PHMM* pipeline developed by Hou et al. [26] evolved from the *OVA* pipeline by introducing a *HMM*-based HCP detection approach to the reference-offspring pairwise-comparison scheme. For each reference chromosome, a crossover can be directly inferred if, within a 1Mb sliding window, HCPs can be identified in over 60% of the reference-offspring pairs. Detailed description of the pipeline can be found in the original paper [26]. Source code of a series of C++ programs and perl scripts for implementing the *PHMM* pipeline are publicly available. To facilitate the comparison analysis in this study, we directly applied the C++ programs for crossover identification but rewrote the perl scripts in R language (without changing the original algorithm) for the inference of consensus haplotypes.

### Maize microspore sequencing dataset

The maize microspore sequencing dataset was generated by Li et al [29]. A total of 96 (24 × 4) microspores from 24 tetrads were isolated from F1 hybrid individuals of a cross between two inbred lines (SK and ZHENG58), and were sequenced at ~1.4× depth coverage. Parents of the F1 hybrid were also sequenced at up to 8× (SK) and 15.7× (ZHENG58) genome coverage depth, respectively. With a stringent filtering process, a total of 599,154 high-quality SNPs were obtained for both parents and the microspores.

### Human sperm sequencing dataset

Single sperm cell sequencing data of 11 sperms from the donor of the HuRef diploid genome were downloaded from the NCBI Sequence Read Archive (SRA) (https://www.ncbi.nlm.nih.gov/sra) with the accession number SRP017516 [28]. Sequences were aligned to the human GRCh37 reference genome using BWA-MEM [32] implemented in the SpeedSeq software [33]. Duplicate-marked, sorted, and indexed BAM files were produced by the SpeedSeq align module, which utilizes SAMBLASTER [34] to mark duplicates and uses Sambamba [35] to sort and index BAM files. For each sperm, the genotypes at 1.95 million heterozygous SNP loci in the HuRef genome were determined using Genome Analysis Toolkit (GATK) [36].

### Data availability

*Hapi* is an R package that is freely available at https://github.com/Jialab-UCR/Hapi

## Acknowledgements

We thank Dr. Jianbing Yan and Dr. Xiang Li at Huazhong Agricultural University for sharing the processed genotype data for the microspores from their maize tetrad study. This work was supported by start-up funding of UCR, UC Academic Senate Regents Faculty Fellowship and Faculty Development Award, and UCR Hellman Fellowship to Zhenyu Jia.

## Author contributions

Z.J. conceived and supervised the study; Z.J., M.L.R., and R.L. designed the study; R.L. and Z.J. developed the pipeline. R.L., H.Q., J.C., S.W., L.Z., and J.W. performed the data analysis. Z.J., R.L., H.Q., J.M.C., and M.L.R. wrote the manuscript with contributions from all authors.

---

**Supplementary Fig. 1:**
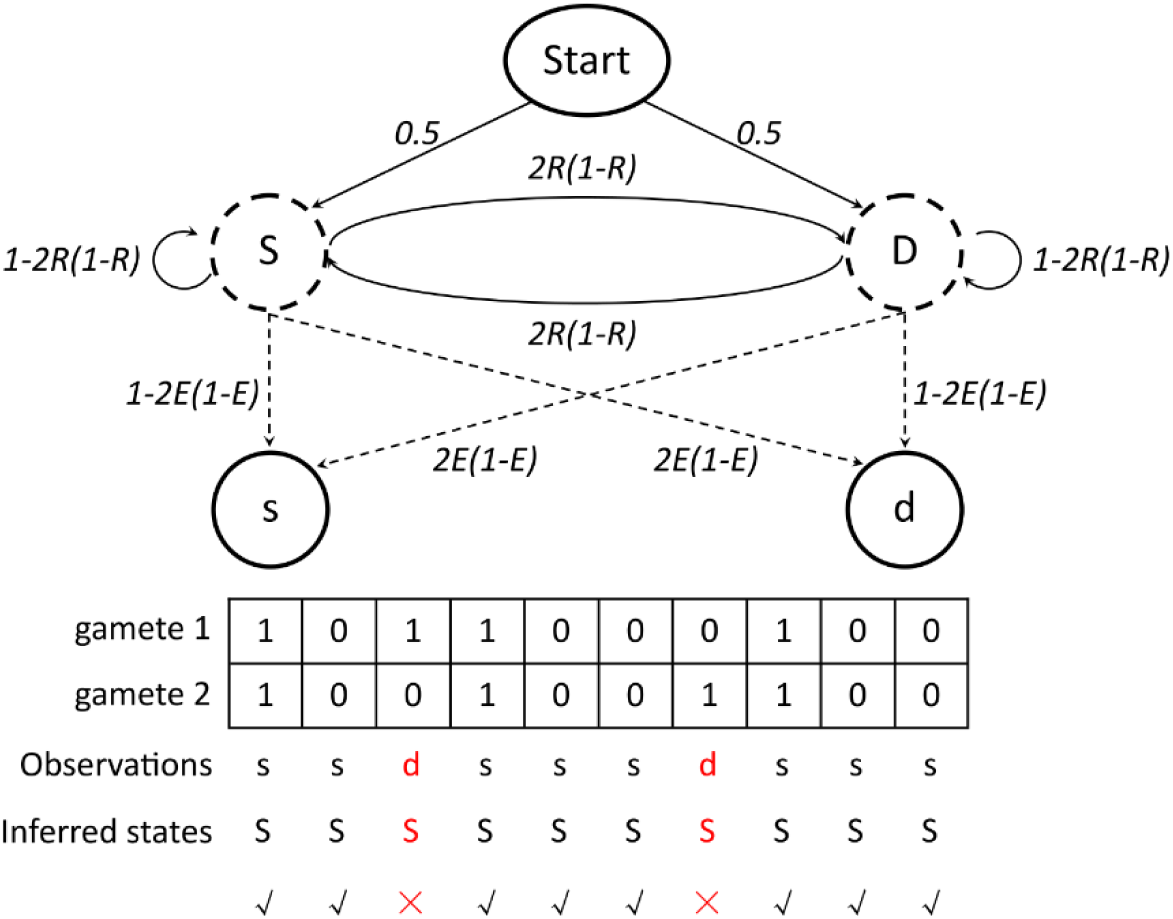
The HMM for detection of hetSNPs with erroneous genotype calls. (*R*: recombination frequency; *E*: genotype error rate; 0 and 1 represent the two alleles that are arbitrarily assigned for a hetSNP locus; × signs in red mark the hetSNPs with potential errors and √ signs in black denote the hetSNPs with correct genotype calls.)

**Supplementary Fig. 2:**
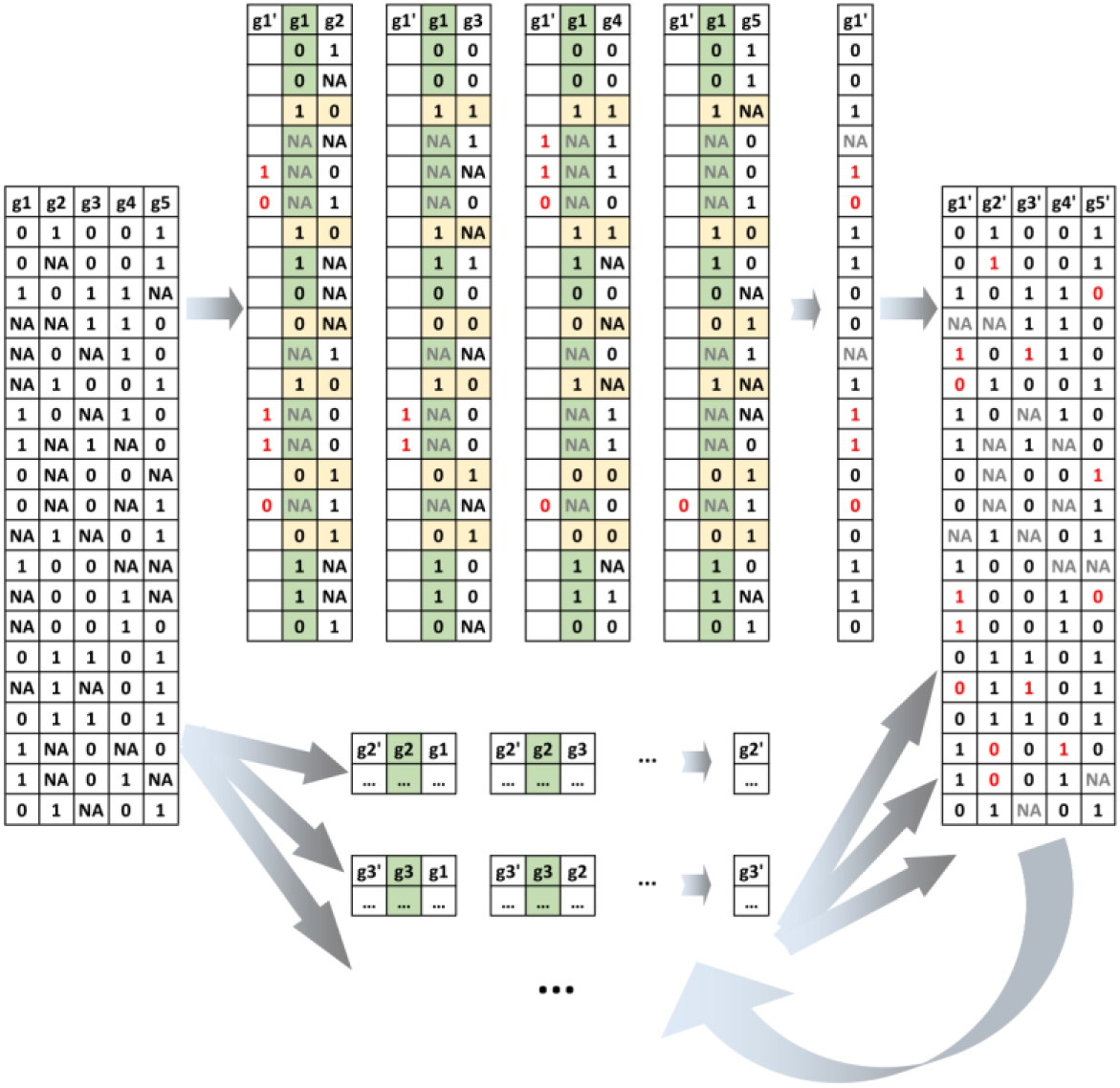
Iterative imputation of missing genotypes in the framework. (g1, …, g5 are the 5 gamete cells with missing genotypes; g1’, …, g5’ are gamete cells after imputation; The gamete with green background is the ‘target gamete’ for imputation; Markers with yellow background are ‘comparator markers’ that immediately around missing regions.)

**Supplementary Fig. 3:**
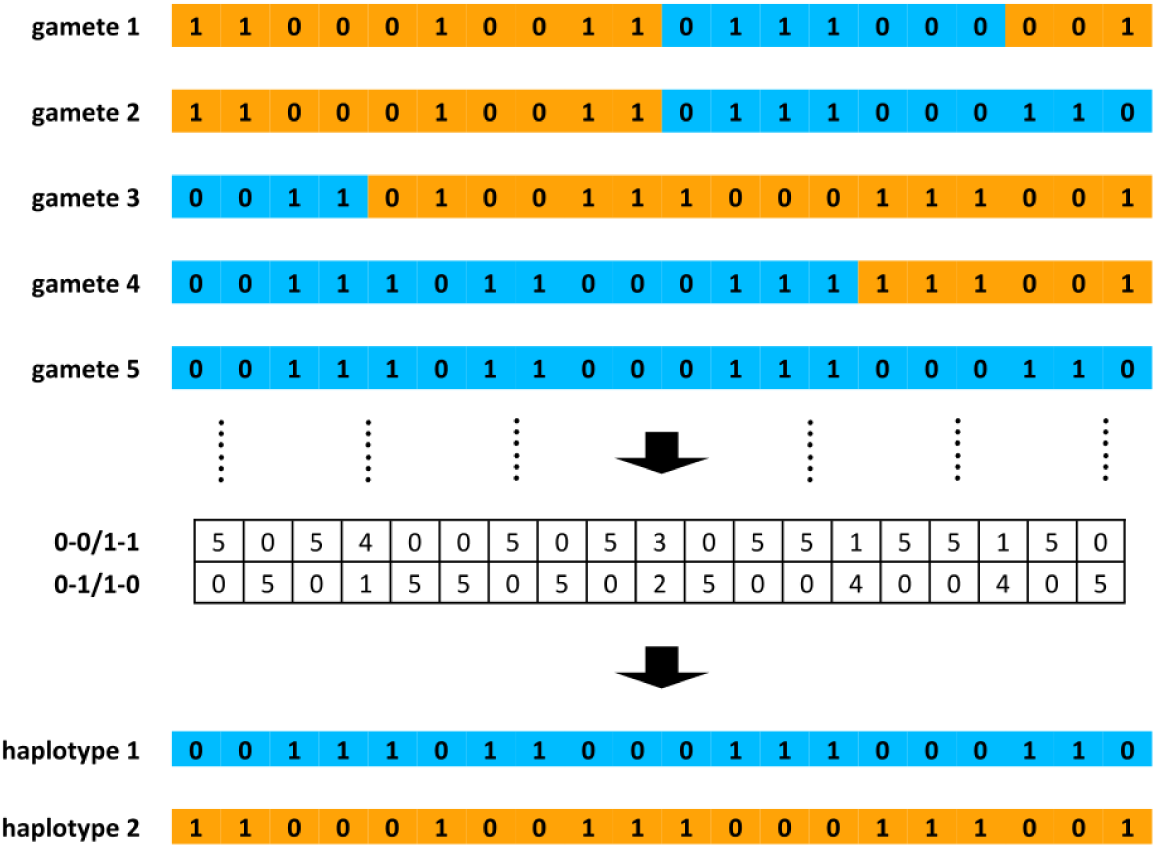
The majority voting strategy for the inference of the draft haplotype. (0 and 1 represent the two alleles that are arbitrarily assigned for a hetSNP locus; The two colors of background indicate the two haplotypes; The table in the middle shows the number of different link types between any two adjacent markers.)

**Supplementary Fig. 4:**
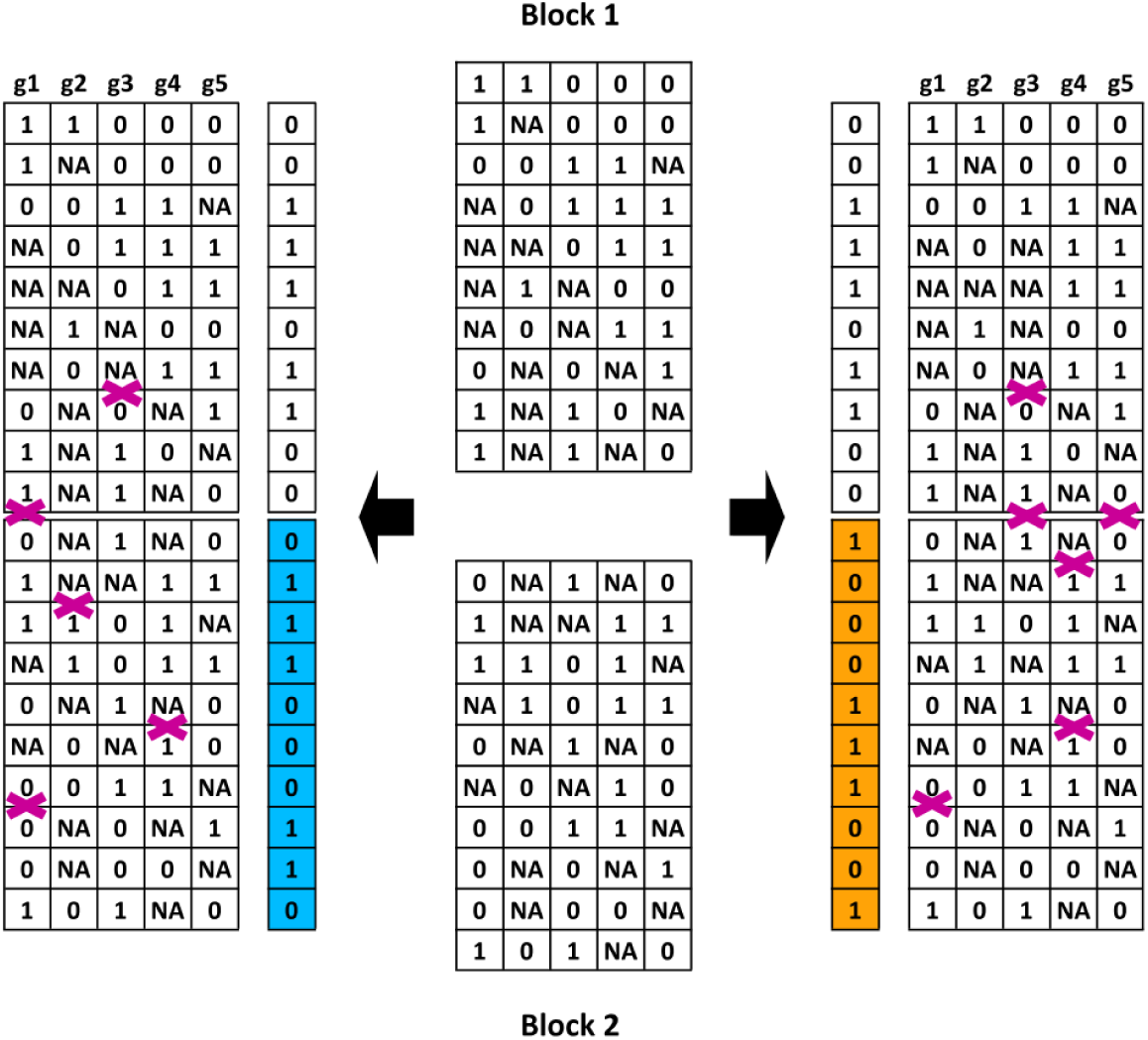
Maximum parsimony of recombination (MPR) principle for proofreading the draft haplotypes. The framework is divided into two blocks (Block 1 and Block 2) at the position under proofreading. There are two possible haplotype combinations between these two blocks. The number of crossovers is counted given each of the two haplotypes and the one generating less crossovers is determined as the true haplotype based upon MPR. (0 and 1 represent the two alleles that are arbitrarily assigned for a hetSNP locus; The two colors of background indicate the two reciprocal haplotypes for Block 2; Purple cross signs denote the positions of crossovers.)

**Supplementary Fig. 5:**
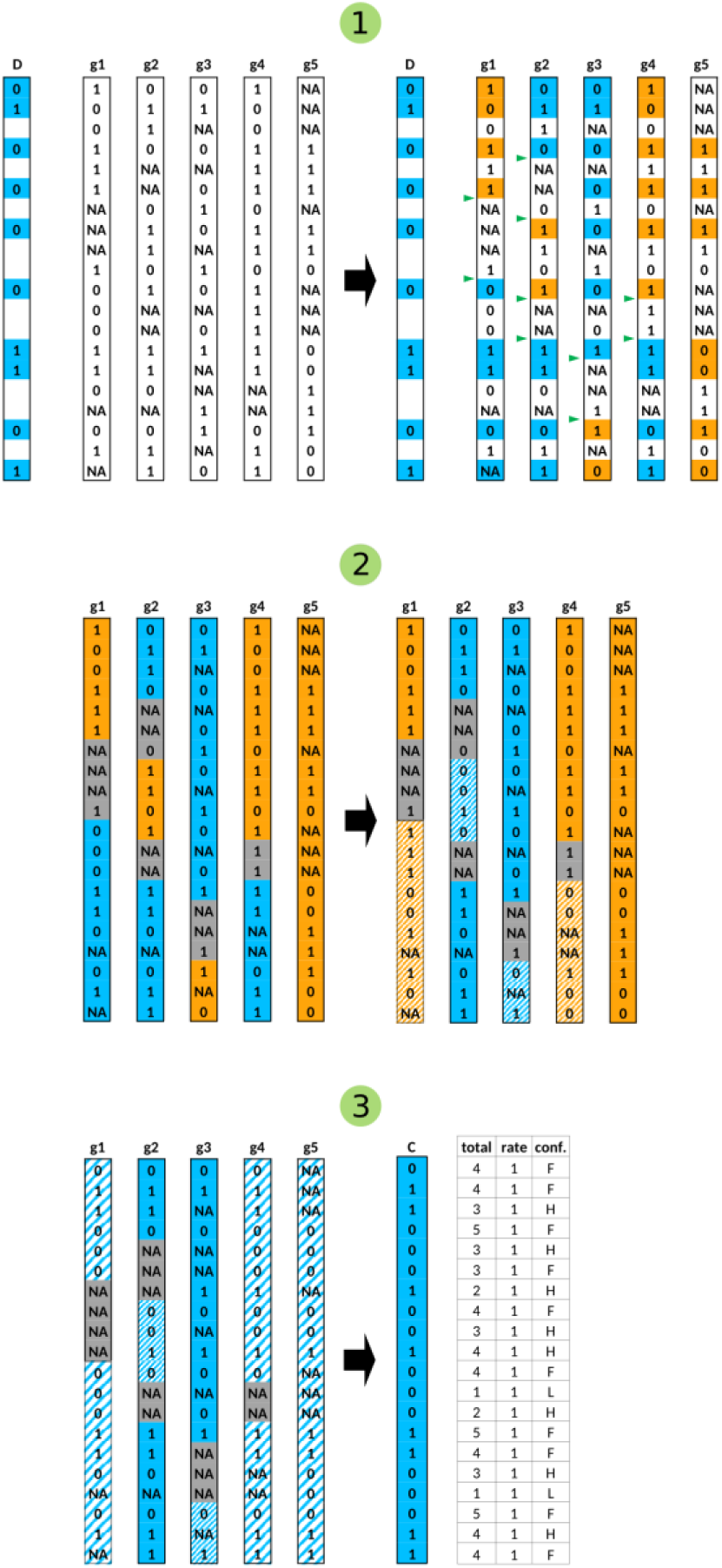
High-resolution consensus haplotype assembly. (g1, …, g5 are the 5 gamete cells; 0 and 1 represent the two alleles that are arbitrarily assigned for a hetSNP locus; D represents draft haplotype and C represents consensus haplotype; In the table of consensus haplotype, ‘total’ is the total number of cells with observed genotypes at a locus, ‘rate’ is the ratio of cells supporting the haplotype, and ‘conf.’ is the confidence of haplotype phasing for each hetSNP, *i.e.*, F indicates that the hetSNP is in the framework, L denotes low-confident phasing, whereas H represents high-confident phasing that is supported by at least 2 gametes and the ratio of cells supporting the haplotype is greater than 0.6; Green triangles mark the HCPs inferred by comparing genotypes in each gamete cell with those in the draft haplotype. The two colors of background indicate the two haplotypes in each gamete, and regions with grey background are unsolved haplotype regions; The regions with slash represent the haplotypes after flipping.)

**Supplementary Fig. 6:**
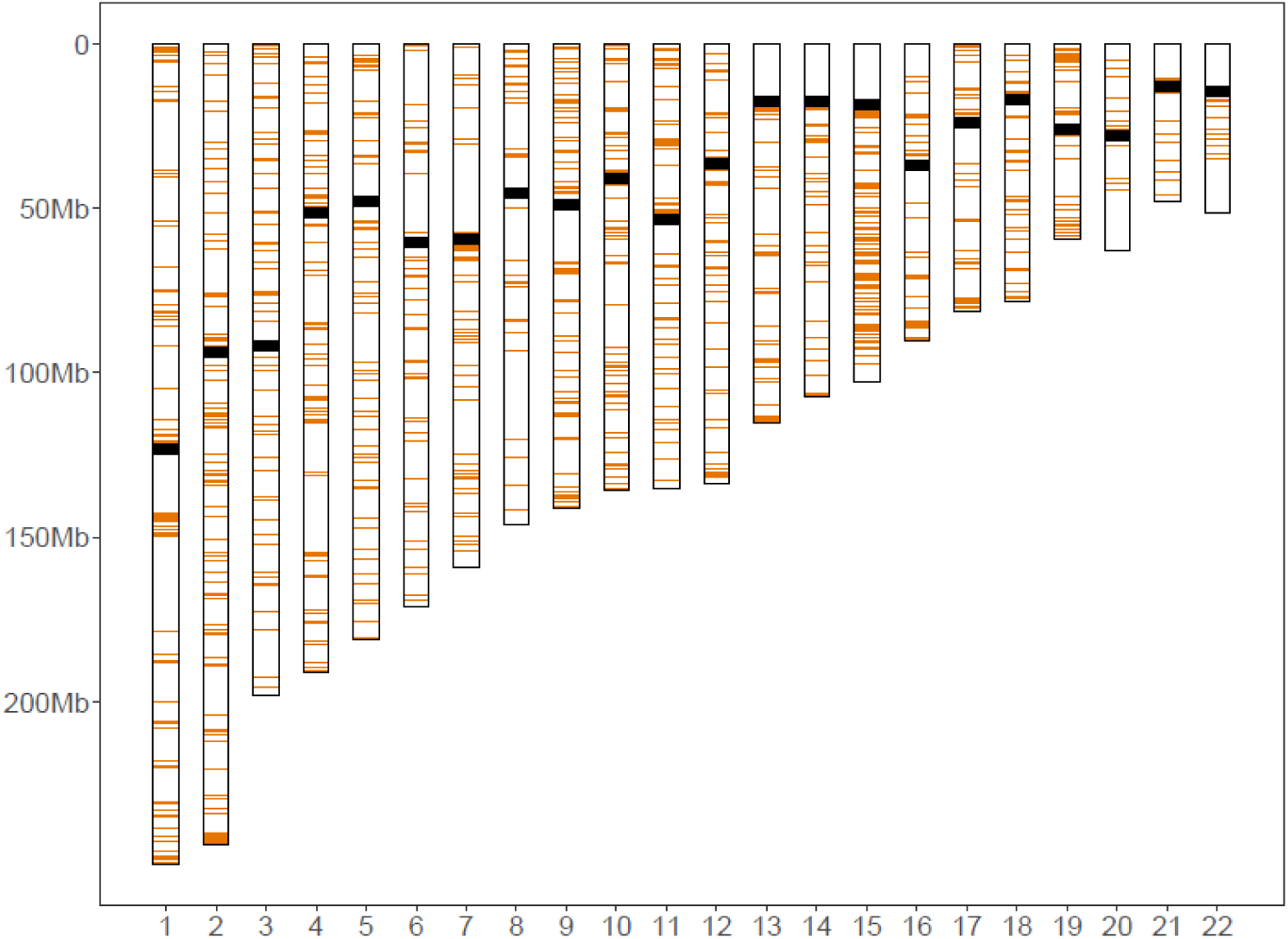
Distribution of phased hetSNPs that disagree between *Hapi* and the suggested haplotypes.

**Supplementary Fig. 7:**
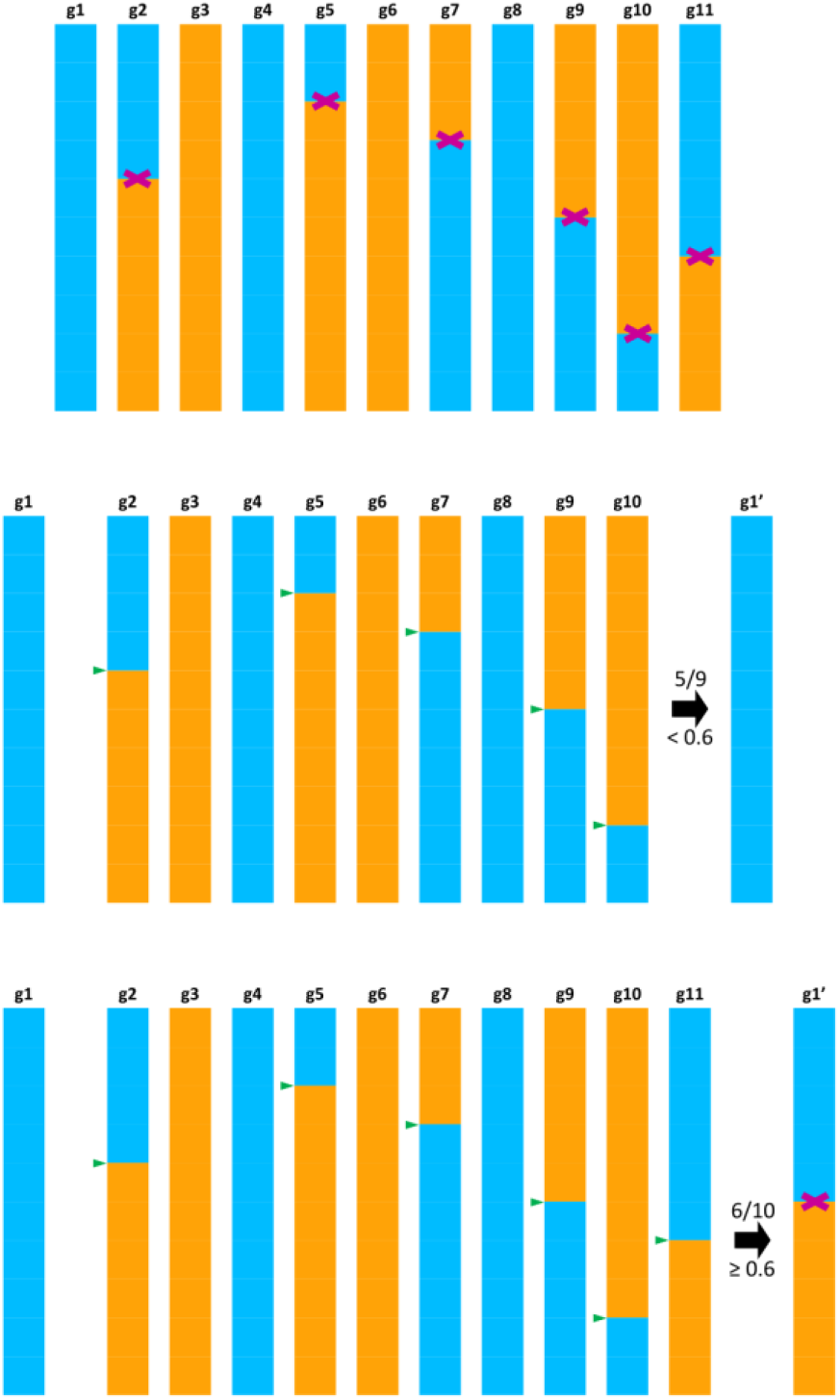
An example of incorrect assignment of crossovers by *PHMM* when multiple crossovers occur in a small region that is less than 1M. (g1, …, g11 are the 11 gamete cells; g1’ is gamete 1 with deduced crossovers via pairwise comparisons with all other gametes; Purple cross signs denote the positions of crossovers and green triangles mark the HCPs inferred by the reference-offspring assay.)

**Supplementary Fig. 8:**
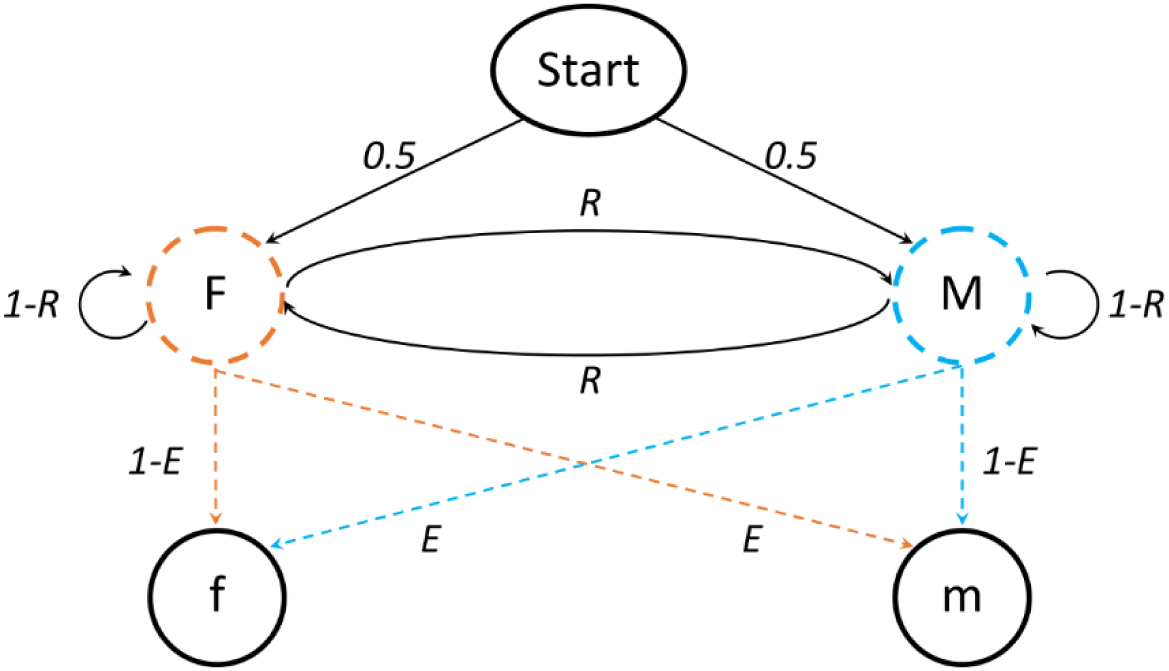
HMM for identification of crossovers. The HMM consists of two observations (f and m), and two hidden states (F and M), representing the paternal and maternal haplotypes, respectively. The initial probabilities of the two states are 0.5. Given the F (or M) state, the emission probability of observing the f (or m) haplotype is *1*-*E* and observing the complementary haplotype m (or f) is *E*, respectively, where *E* is genotype error rate. The transition probabilities from one state to itself is *1*-*R*, and to the other state is *R*, where *R* is recombination frequency. A sequence of hidden states for the ‘chained’ markers can be inferred by Viterbi’s algorithm and crossover positions are determined where state swaps occur.

**Supplementary Fig. 9:**
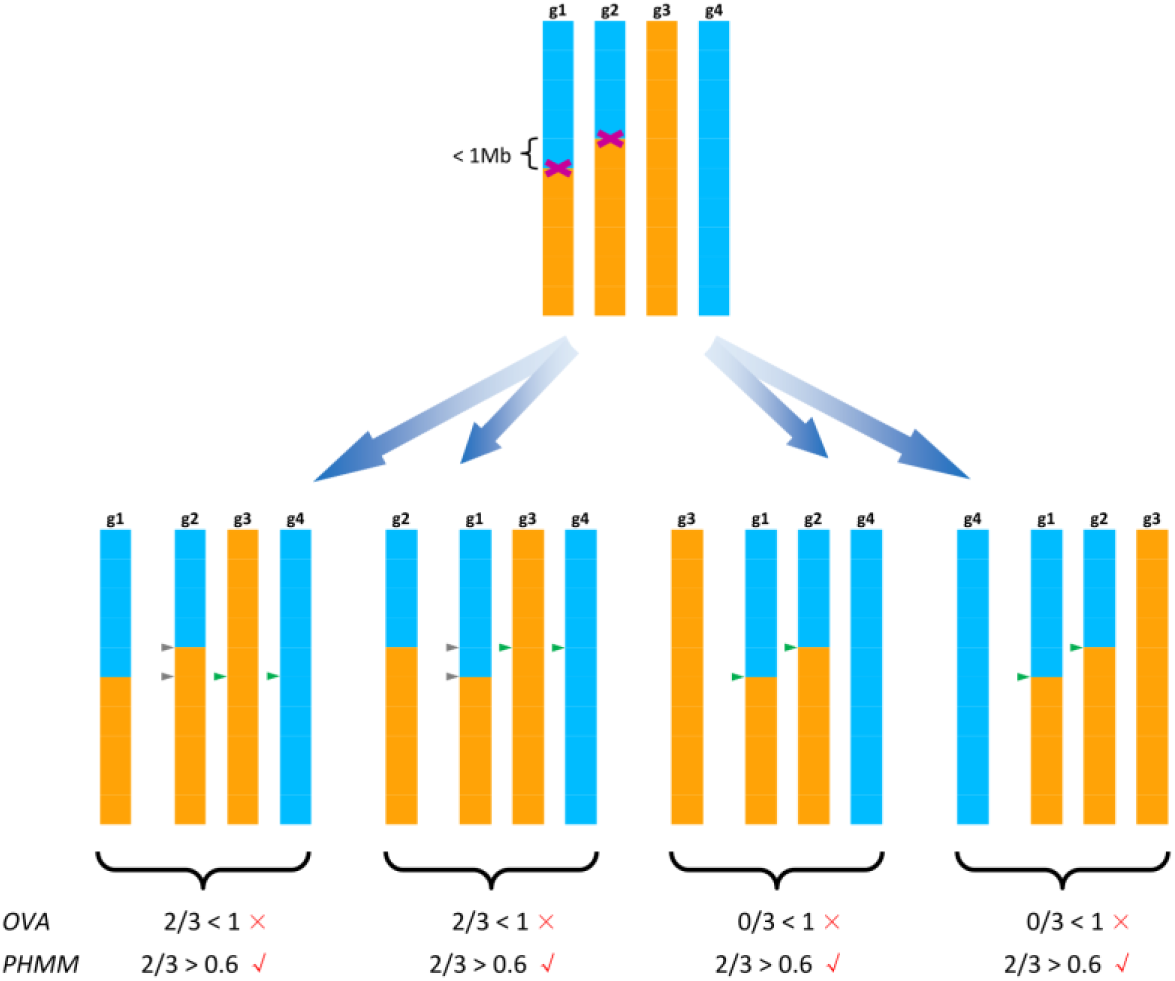
An example of deficiencies for *OVA* and *PHMM* in haplotype phasing when 4 gametes are analyzed and two crossovers (cross signs in purple) occurred in two gametes within 1M chromosomal region. Multiple HCPs within a 1 Mb region inferred from a reference-offspring pair are filtered out by both methods (triangles in gray). *OVA* detects HCPs (triangles in green) with overlapping hetSNPs and uses a threshold (rate of support from the reference-offspring assay) of 1 to assign a crossover to the reference chromosome. *PHMM* searches HCPs (triangles in green) in a 1Mb sliding window and uses a cutoff of 0.6 to determine the crossover on the reference chromosome. In this case, OVA identified no crossover (4 X signs in red), while 4 crossovers (4 √ signs in red) have been detected by *PHMM* with each gamete bearing one crossover.

## References

1. Gregoriussen M, Maria A, Bohr HG: A novel model on DST-induced transplantation tolerance by the transfer of self-specific donor tTregs to a haplotype-matched organ recipient. Frontiers in immunology 2017, 8:9.

2. Glusman G, Cox HC, Roach JC: Whole-genome haplotyping approaches and genomic medicine. Genome medicine 2014, 6:73.

3. McCarthy S, Das S, Kretzschmar W, Delaneau O, Wood AR, Teumer A, Kang HM, Fuchsberger C, Danecek P, Sharp K: A reference panel of 64,976 haplotypes for genotype imputation. Nature genetics 2016, 48:1279.

4. Huang J, Howie B, McCarthy S, Memari Y, Walter K, Min JL, Danecek P, Malerba G, Trabetti E, Zheng H-F: Improved imputation of low-frequency and rare variants using the UK10K haplotype reference panel. Nature communications 2015, 6:8111.

5. Lambert J-C, Grenier-Boley B, Harold D, Zelenika D, Chouraki V, Kamatani Y, Sleegers K, Ikram M, Hiltunen M, Reitz C: Genome-wide haplotype association study identifies the FRMD4A gene as a risk locus for Alzheimer’s disease. Molecular psychiatry 2013, 18:461.

6. Trégouët D-A, König IR, Erdmann J, Munteanu A, Braund PS, Hall AS, Großhennig A, Linsel-Nitschke P, Perret C, DeSuremain M: Genome-wide haplotype association study identifies the SLC22A3-LPAL2-LPA gene cluster as a risk locus for coronary artery disease. Nature genetics 2009, 41:283.

7. Yang J, Ferreira T, Morris AP, Medland SE, Madden PA, Heath AC, Martin NG, Montgomery GW, Weedon MN, Loos RJ: Conditional and joint multiple-SNP analysis of GWAS summary statistics identifies additional variants influencing complex traits. Nature genetics 2012, 44:369.

8. de Bakker PI, Yelensky R, Pe’er I, Gabriel SB, Daly MJ, Altshuler D: Efficiency and power in genetic association studies. Nature genetics 2005, 37:1217.

9. Browning SR, Browning BL: Rapid and accurate haplotype phasing and missing-data inference for whole-genome association studies by use of localized haplotype clustering. The American Journal of Human Genetics 2007, 81:1084–1097.

10. Li Y, Willer CJ, Ding J, Scheet P, Abecasis GR: MaCH: using sequence and genotype data to estimate haplotypes and unobserved genotypes. Genetic epidemiology 2010, 34:816–834.

11. Loh P-R, Danecek P, Palamara PF, Fuchsberger C, Reshef YA, Finucane HK, Schoenherr S, Forer L, McCarthy S, Abecasis GR: Reference-based phasing using the Haplotype Reference Consortium panel. Nature genetics 2016, 48:1443.

12. O’Connell J, Sharp K, Shrine N, Wain L, Hall I, Tobin M, Zagury J-F, Delaneau O, Marchini J: Haplotype estimation for biobank-scale data sets. Nature genetics 2016, 48:817.

13. Stephens M, Scheet P: Accounting for decay of linkage disequilibrium in haplotype inference and missing-data imputation. The American Journal of Human Genetics 2005, 76:449–462.

14. Stephens M, Smith NJ, Donnelly P: A new statistical method for haplotype reconstruction from population data. The American Journal of Human Genetics 2001, 68:978–989.

15. Scheet P, Stephens M: A fast and flexible statistical model for large-scale population genotype data: applications to inferring missing genotypes and haplotypic phase. The American Journal of Human Genetics 2006, 78:629–644.

16. Howie BN, Donnelly P, Marchini J: A flexible and accurate genotype imputation method for the next generation of genome-wide association studies. PLoS genetics 2009, 5:e1000529.

17. Ma L, Xiao Y, Huang H, Wang Q, Rao W, Feng Y, Zhang K, Song Q: Direct determination of molecular haplotypes by chromosome microdissection. Nature methods 2010, 7:299.

18. Yang H, Chen X, Wong WH: Completely phased genome sequencing through chromosome sorting. Proceedings of the National Academy of Sciences 2011, 108:12-17.

19. Fan HC, Wang J, Potanina A, Quake SR: Whole-genome molecular haplotyping of single cells. Nature biotechnology 2011,29:51.

20. Peters BA, Kermani BG, Sparks AB, Alferov O, Hong P, Alexeev A, Jiang Y, Dahl F, Tang YT, Haas J: Accurate whole-genome sequencing and haplotyping from 10 to 20 human cells. Nature 2012, 487:190.

21. Selvaraj S, Dixon JR, Bansal V, Ren B: Whole-genome haplotype reconstruction using proximity-ligation and shotgun sequencing. Nature biotechnology 2013, 31:1111.

22. Edge P, Bafna V, Bansal V: HapCUT2: robust and accurate haplotype assembly for diverse sequencing technologies. Genome research 2017, 27:801–812.

23. Kitzman JO, MacKenzie AP, Adey A, Hiatt JB, Patwardhan RP, Sudmant PH, Ng SB, Alkan C, Qiu R, Eichler EE: Haplotype-resolved genome sequencing of a Gujarati Indian individual. Nature biotechnology 2010, 29:59.

24. Porubsky D, Garg S, Sanders AD, Korbel JO, Guryev V, Lansdorp PM, Marschall T: Dense and accurate whole-chromosome haplotyping of individual genomes. Nature Communications 2017, 8:1293.

25. Porubský D, Sanders AD, van Wietmarschen N, Falconer E, Hills M, Spierings DC, Bevova MR, Guryev V, Lansdorp PM: Direct chromosome-length haplotyping by single-cell sequencing. Genome research 2016, 26:1565–1574.

26. Hou Y, Fan W, Yan L, Li R, Lian Y, Huang J, Li J, Xu L, Tang F, Xie XS: Genome analyses of single human oocytes. Cell 2013, 155:1492–1506.

27. Lu S, Zong C, Fan W, Yang M, Li J, Chapman AR, Zhu P, Hu X, Xu L, Yan L: Probing meiotic recombination and aneuploidy of single sperm cells by whole-genome sequencing. Science 2012, 338:1627–1630.

28. Kirkness EF, Grindberg RV, Yee-Greenbaum J, Marshall CR, Scherer SW, Lasken RS, Venter JC: Sequencing of isolated sperm cells for direct haplotyping of a human genome. Genome research 2013, 23:826–832.

29. Li X, Li L, Yan J: Dissecting meiotic recombination based on tetrad analysis by single-microspore sequencing in maize. Nature communications 2015, 6:6648.

30. Viterbi A: Error bounds for convolutional codes and an asymptotically optimum decoding algorithm. IEEE transactions on Information Theory 1967, 13:260–269.

31. Xie W, Feng Q, Yu H, Huang X, Zhao Q, Xing Y, Yu S, Han B, Zhang Q: Parent-independent genotyping for constructing an ultrahigh-density linkage map based on population sequencing. Proceedings of the National Academy of Sciences 2010, 107:10578–10583.

32. Li H, Durbin R: Fast and accurate short read alignment with Burrows-Wheeler transform. Bioinformatics 2009, 25:1754–1760.

33. Chiang C, Layer RM, Faust GG, Lindberg MR, Rose DB, Garrison EP, Marth GT, Quinlan AR, Hall IM: SpeedSeq: ultra-fast personal genome analysis and interpretation. Nature methods 2015, 12:966.

34. Faust GG, Hall IM: SAMBLASTER: fast duplicate marking and structural variant read extraction. Bioinformatics 2014, 30:2503–2505.

35. Tarasov A, Vilella AJ, Cuppen E, Nijman IJ, Prins P: Sambamba: fast processing of NGS alignment formats. Bioinformatics 2015, 31:2032–2034.

36. DePristo MA, Banks E, Poplin R, Garimella KV, Maguire JR, Hartl C, Philippakis AA, Del Angel G, Rivas MA, Hanna M: A framework for variation discovery and genotyping using next-generation DNA sequencing data. Nature genetics 2011,43:491.

